# Mammalian SWI/SNF chromatin remodeler is essential for reductional meiosis in males

**DOI:** 10.1101/2020.04.28.066647

**Authors:** Debashish U. Menon, Terry Magnuson

## Abstract

BRG1, a catalytic subunit of the mammalian SWI/SNF nucleosome remodeler is essential for male meiosis^1^. In addition to BRG1, multiple subunits (~10-14) some of which are mutually exclusive, constitute biochemically distinct SWI/SNF subcomplexes, whose functions in gametogenesis remain unknown. Here, we identify a role for the PBAF (Polybromo <u>- B</u>rg1 <u>A</u>ssociated <u>F</u>actor) complex in the regulation of meiotic cell division. The germ cell-specific depletion of PBAF specific subunit, ARID2 resulted in a metaphase-I arrest. *Arid2^cKO^* metaphase-I spermatocytes displayed defects in chromosome organization and spindle assembly. Additionally, mutant centromeres were devoid of Polo-like kinase1 (PLK1), a known regulator of the spindle assembly checkpoint (SAC)^2^. The loss of PLK1 coincided with an abnormal chromosome-wide expansion of centromeric chromatin modifications such as Histone H3 threonine3 phosphorylation (H3T3P) and Histone H2A threonine120 phosphorylation (H2AT120P) that are critical for chromosome segregation^3,4^. Consistent with the known role of these histone modifications in chromosome passenger complex (CPC) recruitment, *Arid2^cKO^* metaphase-I chromosomes display defects in CPC association. We propose that ARID2 facilitates metaphase-I exit by regulating spindle assembly and centromeric chromatin.

## Main text

We previously demonstrated a role for the mammalian SWI/SNF chromatin remodeling complex in meiotic progression by conditionally ablating its ATPase subunit, BRG1, in the testis^1,5^. Given that SWI/SNF complexes are biochemically heterogenous^6,7^, it is possible that certain subcomplexes might selectively govern distinct spermatogenic processes. To address this possibility, we examined existing RNA-seq data generated from purified spermatogenic cell populations^8^ and monitored the temporal expression profiles of SWI/SNF subunits that define two subcomplexes, namely, BAF (<u>B</u>rg1 <u>A</u>ssociated <u>F</u>actor) and PBAF (Polybromo-BAF) (Fig. 1a). Of particular interest were the ARID (<u>AT</u>-<u>r</u>ich interaction <u>d</u>omain) containing SWI/SNF subunits, known to bind DNA with loose specificity^9^. Among the subunits examined, the mRNA expression of PBAF DNA binding subunit^10^, *Arid2,* drew our attention as it peaked at pachynema (Fig. 1a). More recent single cell RNA seq data also identified *Arid2* as an early pachytene marker^11^, suggesting a role in meiosis-I. We therefore performed immunostaining on testes cryosections to profile ARID2 abundance in prophase-I spermatocytes staged by γH2Ax. Consistent with its mRNA profile, ARID2 went from being undetectable to expressed weakly, during the transition from pre-leptonema to zygonema, but appeared elevated at pachynema which is distinguished by the sex body^12^. Additionally, ARID2 was also detected in round spermatids, enriched proximally to chromocenters (Fig. 1b). To investigate its role in meiosis we conditionally knocked out *Arid2* using the spermatogonia specific *Stra8-Cre* transgene^13^, resulting in significant depletion of ARID2 and smaller testes in *Arid2^cKO^* relative to *Arid2^WT^* (Extended Data Fig. 1a, Fig. 1c). A histological examination revealed a striking loss of haploid spermiogenic cell populations and absence of mature spermatozoa in *Arid2^cKO^* relative to *Arid2^WT^* adult seminiferous tubules and cauda epididymides respectively (Fig. 1d, Extended Data Fig. 1b). The *Arid2^cKO^* testis were characterized by an accumulation of stage XII tubules which is indicative of a meiotic arrest (Fig. 1d). While meiotic prophase-I appeared unhindered, we noticed a striking accumulation of metaphase - I spermatocytes in *Arid2^cKO^* testis (Fig. 1d, e, Extended Data Fig. 1c). The near absence of secondary spermatocytes and spermiogenic cells in *Arid2^cKO^* testis, indicate that ARID2 is essential for meiotic cell division.

**Figure 1.**
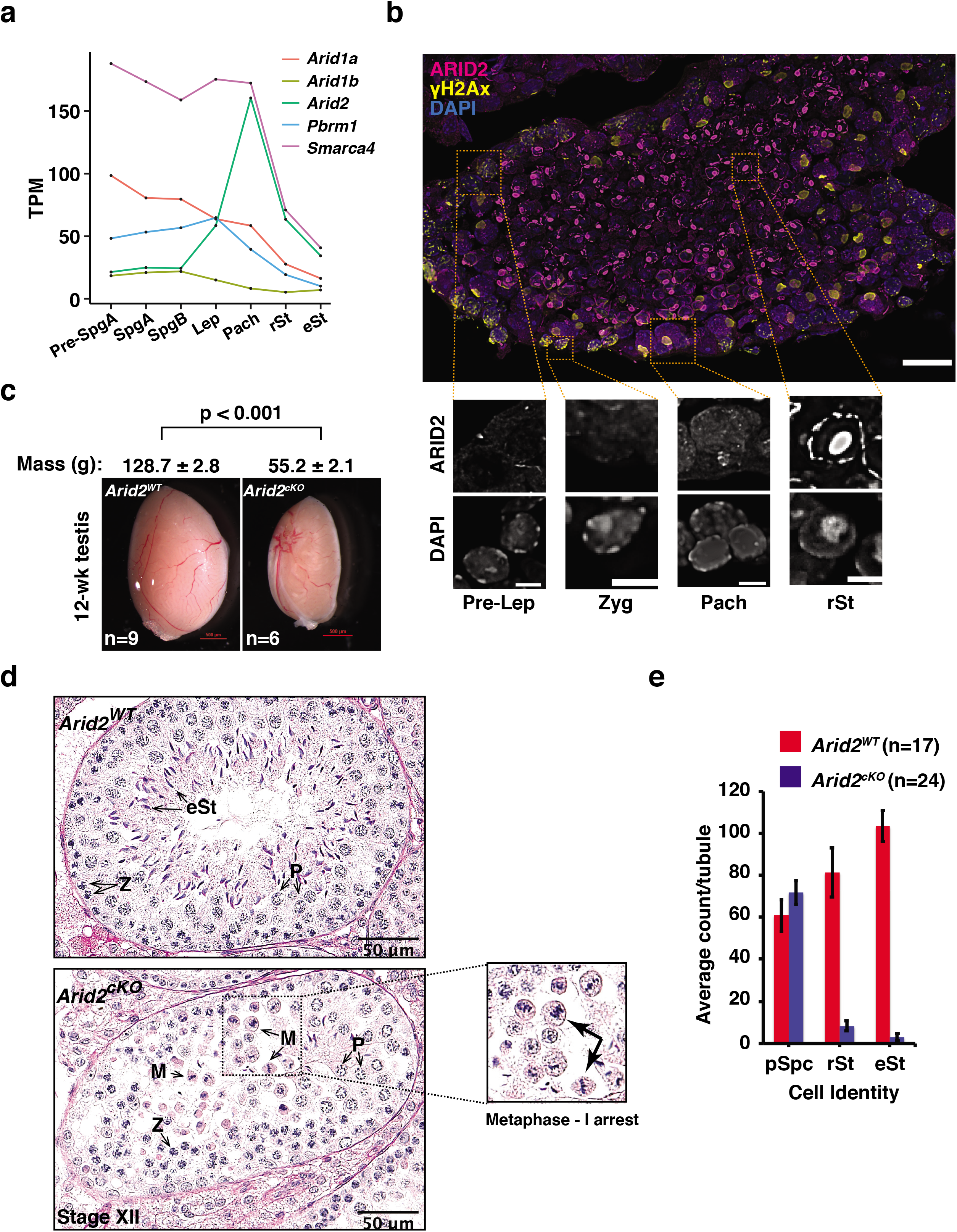
The loss of ARID2 results in a late meiosis-I arrest. (a) mRNA expression profile of SWI/SNF subunits in purified germ cell populations^8^. (b) Adult testis cryosection immunolabelled for ARID2 (magenta), gH2Ax (yellow) and counterstained with DAPI (blue), scale bar: 20 μm. Magnified views of representative cell types are displayed below, scale bar: 5 μm. Magnification:40x. (c) *Arid2^WT^* and *Arid2^cKO^* testis mounts. Average testis mass (g) and standard error obtained from multiple measurements (n) are indicated. p-value calculated using students t-test. Scale bar: 500 μm. (d) 12-week-old *Arid2^WT^ and Arid2^cKO^* PAS-stained testes sections. Magnified view of *Arid2^cKO^* metaphase spermatocytes are displayed. Scale bar: 50 μm, magnification: 40x. (e) Proportions of prophase-I spermatocytes (pSpc) and spermiogenic cell populations quantified from 12-week-old *Arid2^WT^* and *Arid2^cKO^* PAS-stained testes sections. Number of tubules (n) examined for each genotype are indicated. (a,b,d) Spg: Spermatogonia, Lep: Leptotene spermatocyte, Pach/P: Pachytene spermatocyte, Zyg/Z: Zygotene spermatocyte, M: Metaphase-I spermatocyte, rSt: round spermatid and eSt: elongated spermatid.

Consistent with a putative role in cell division, we were able to detect abundant expression of both ARID2 and BRG1 in metaphase-I spermatocytes by immunofluorescence (IF). They appeared localized peripherally to the metaphase plate displaying limited chromosomal association (Extended Data Fig. 1d, top row). While transitioning to anaphase-I, both ARID2 and BRG1 were detected at the spindle mid-zone while maintaining their peripheral association with segregating chromosomes (Extended Data Fig. 1d, bottom row). The primarily cytosolic distribution of SWI/SNF combined with its narrow localization to chromosomes might suggest a preferential association with the spindle and or kinetochores. Based on the mutant phenotype and spatial distribution of PBAF subunits in late meiosis-I spermatocytes, we hypothesized that ARID2 regulates processes essential for chromosome segregation. These include chromosome condensation, alignment and microtubule-kinetochore attachment.

We began by monitoring phosphorylated histone H3 on serine 10 (H3S10P), a known marker of chromosome condensation and β-Tubulin which constitutes the spindle apparatus, in Arid*2^WT^* and Arid*2^cKO^* seminiferous tubules. While H3S10P levels between *Arid2^WT^* and *Arid2^cKO^* metaphase-I spermatocytes looked similar, certain mutant metaphase-I chromosomes appeared rounded and failed to congress at the metaphase plate (Fig. 2a, 3^rd^ row). These defects prompted us to examine meiotic metaphase spreads where we observed no signs of pairing abnormalities or aneuploidy in *Arid2^cKO^* spermatocytes (Extended Fig. 2a). More strikingly, 52 % of the scored metaphase-I spermatocytes from *Arid2^cKO^* tubules lacked a normal spindle (Fig. 2a, 3^rd^ row). Metaphase-I spermatocytes that lacked spindle were demonstrably deficient in ARID2 relative to those that displayed spindle in *Arid2^cKO^* tubules (Extended Fig. 2b). The latter therefore represent cells that might have undergone inefficient knockout and were considered internal controls. Therefore, *Arid2^cKO^* metaphase-I spermatocytes feature aberrant spindle assembly and abnormal chromosome organization.

**Figure 2.**
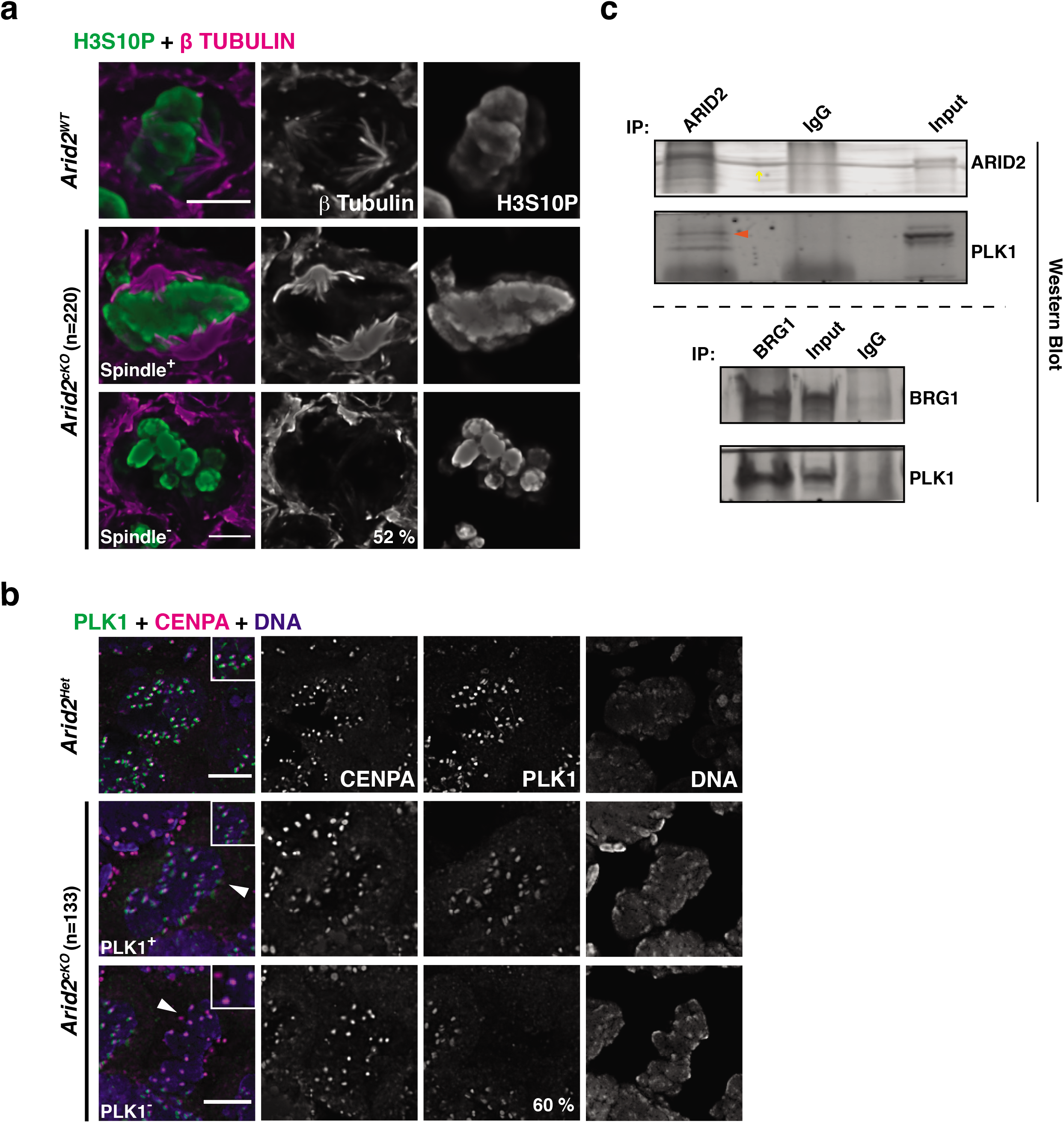
ARID2 influences spindle assembly and PLK1 association with kinetochore. (a,b) Metaphase-I spermatocytes from control and *Arid2^cKO^* testes cryosections immunolabelled for (a) H3S10P (green) and β-Tubulin (magenta), (b) PLK1 (green), CENPA (magenta) and counterstained with DAPI (blue). Arrowheads denote cell of interest and panel insets highlight centromeres. (a,b) Scale bars: 5 μm, magnification:100x. Internal control (Spindle^+^/PLK1^+^) and mutant metaphase-I spermatocytes (Spindle7PLK1^) were imaged from same section. Total number of metaphase-I spermatocytes scored (n) from *Arid2^cKO^* seminiferous tubules and proportion (%) of mutant cells are indicated. (c) ARID2 (top) and BRG1 (bottom) Co-IP. Red arrowhead indicates interacting protein, yellow arrow denotes non-specific streak.

Normally, a failure to establish microtubule – kinetochore attachments would activate the spindle assembly checkpoint (SAC) which delays metaphase exit^14,15^. Given that *Arid2^cKO^* spermatocytes are deficient in microtubules we were curious to determine the status of SAC proteins many of which associate with kinetochores. For this purpose, we chose Polo-like kinase1 (PLK1), a well-known SAC regulator^2,16^ that also influences spindle assembly^17–22^. In control *(Arid2^Het^)* metaphase-I spermatocytes, PLK1 foci were seen overlapping with Centromere protein A (CENPA), a known histone H3 variant and component of functional centromeres^23^ (Fig. 2b). In contrast, 60 % of the scored metaphase-I spermatocytes in *Arid2^cKO^* tubules lacked centromeric PLK1 (PLK1^-^, Fig. 2b). Those that still displayed centromeric PLK1 (PLK1^+^) appeared to express ARID2 at almost normal levels, indicating that they were internal controls (Extended Fig. 2c). Since PLK1 is known to be expressed prior to metaphase-I^24^, we monitored PLK1 in diplotene spermatocytes where desynapsing axial elements are marked by HORMAD1 ^25^. Here, PLK1 foci appeared stable in ARID2 deficient cells (Extended Fig. 2d). Therefore, the destabilization of PLK1 was specific to *Arid2^cKO^* metaphase-I spermatocytes. Interestingly, PLK1 was detected in ARID2 and BRG1 co-immunoprecipitations (Co-IP) with nuclear lysates from spermatogenic cells harvested at 3-weeks post-partum (Fig. 2c). This suggests that PBAF might facilitate the recruitment of PLK1 to centromeres.

Given that PLK1 is known to influence centromeric chromatin during mitosis ^26^, we were curious to determine the impact of ARID2 on histone modifications known to regulate cell division. These include Histone H3 threonine3 phosphorylation (H3T3P) and Histone H2A threonine120 phosphorylation (H2AT120P), that are concomitantly deposited by HASPIN kinase^3,27^ and spindle checkpoint kinase BUB1 ^4^ respectively. Interestingly, perturbation of H3T3P levels have been shown to be detrimental to oocyte maturation in mouse ^27^. Therefore, we first monitored H3T3P in *Arid2^WT^* (normal) metaphase-I spermatocyte squashes. Similar to metaphase-I oocytes, H3T3P overlapped centromeres and was distributed along the inter-chromatid axes (ICA) (Fig. 3a, 1st row). In *Arid2^cKO^* preparations, the absence of centromeric PLK1 was used to differentiate the mutant metaphase-I spermatocytes from internal controls (Fig. 2b, Extended Fig. 2c). H3T3P distribution was limited in internal controls (10%) resembling normal metaphase-I spermatocytes (Fig. 3a, 2nd row). In contrast to normal spermatocytes and internal controls, H3T3P was spread chromosome-wide in majority of mutant metaphase-I spermatocytes (60%), with fewer mutants (30%) displaying low levels of chromatid bound H3T3P (Fig. 3a, row 3,4). We also monitored H3T3P abundance in spermatogenic histone extracts obtained from control and *Arid2^cKO^* testes at time points spanning the appearance of metaphase spermatocytes. By western blot, we failed to notice enhanced H3T3P levels upon the loss of ARID2 (Extended Fig. 3a, top panel). Therefore, the chromosome-wide expansion of H3T3P in mutants is unlikely to occur from increased HASPIN activity.

**Figure 3.**
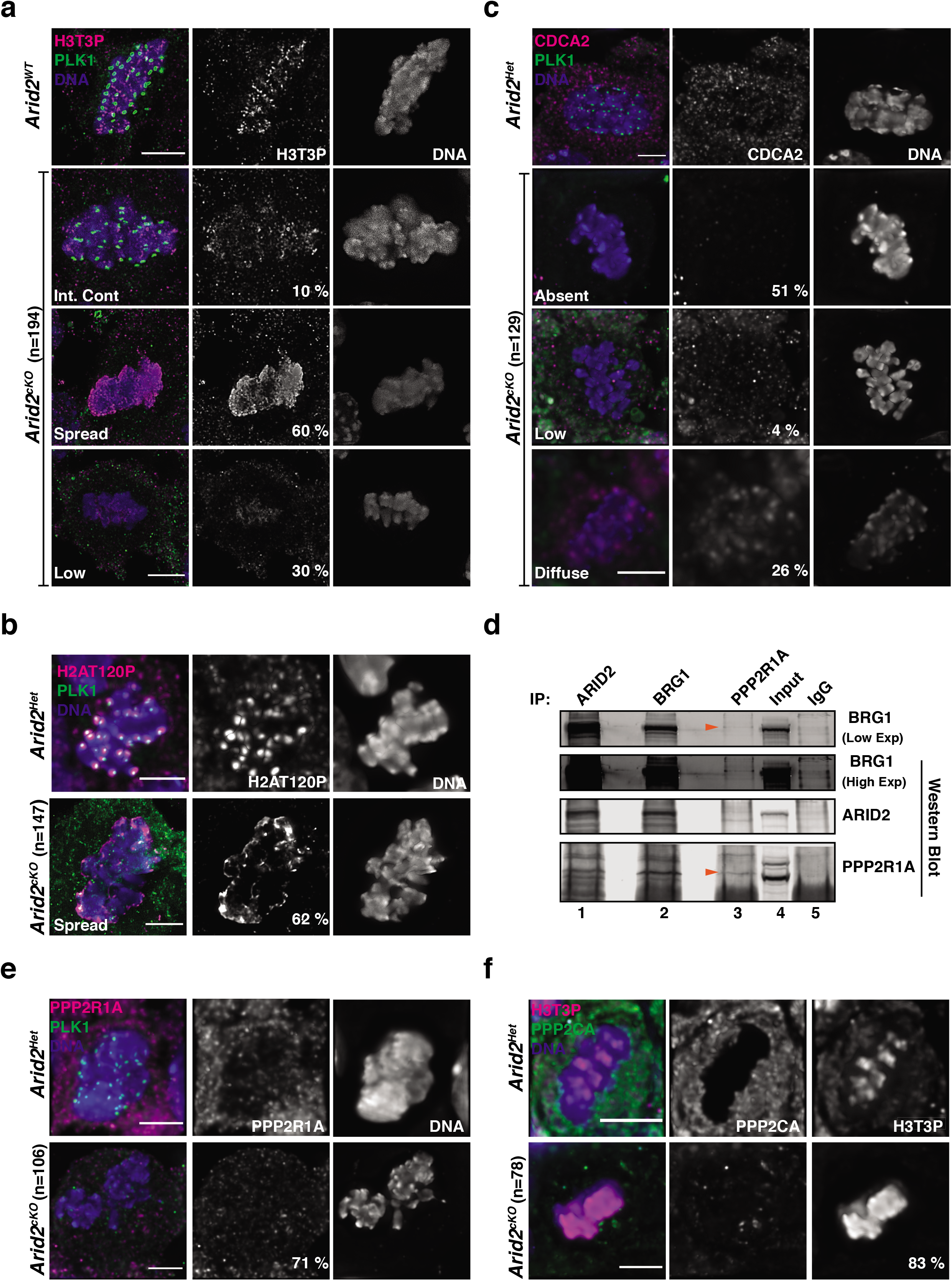
ARID2 regulates centromeric chromatin in metaphase-I spermatocytes. (a-c) Control and *Arid2^cKO^* metaphase-I spermatocyte squashes immunolabelled for PLK1 (green) and (a) H3T3P (magenta), (b) H2AT120P (magenta) and (c) CDCA2 (magenta). DNA stained with DAPI. Scale bar: 5 μm, magnification: 100.8x. (d) ARID2, BRG1 and PPP2R1A Co-IP’s. Red arrowheads indicate interacting proteins. Numbers at bottom denote sample lanes. (e,f) Control and *Arid2^cKO^* metaphase-I spermatocytes immunolabelled for (e) PPP2R1A (magenta) and PLK1 (green), (f) H3T3P (magenta) and PPP2CA (green). Scale bar: 5 μm, magnification: 100.8x. (a-c, e-f) Total number of metaphase-I spermatocytes scored (n) from *Arid2^cKO^* seminiferous tubules and proportion (%) of each abnormal category are indicated.

Next, we determined the impact of ARID2 on H2AT120P. During mitosis, H2AT120P is restricted to centromeres^4,28^. A similar centromeric enrichment of H2A120P was seen in *Arid2^Het^* (control) metaphase-I spermatocytes (Fig. 3a, b, 1^st^ row). In contrast, mutant metaphase-I spermatocytes (62%) exhibited ectopic spreading of H2AT120P (Fig. 3b, 2^nd^ row). The loss of ARID2 did not alter total H2AT120P abundance, indicating normal BUB1 activity in the mutant (Extended Fig. 3b). Therefore, ARID2 is required to ensure the centromeric enrichment of both H3T3P and H2AT120P.

In addition to centromeric chromatin we also determined whether flanking pericentric chromatin, normally marked by H3K9me3^29^ was impacted in the absence of ARID2. Like the centromeric marks, total H3K9me3 abundance was not influenced by the loss of ARID2 (Extended Fig. 3a, bottom panel). However, many mutant metaphase-I spermatocytes from *Arid2^cKO^* testis displayed H3K9me3 spreading (38%) with the remaining exhibiting normal pericentric enrichment (30%), similar to internal controls (26%) and *Arid2^Het^* spermatocytes (Extended Fig. 3c). Therefore, along with centromeric chromatin ARID2 also influences pericentric chromatin organization, albeit at reduced penetrance.

To understand how ARID2 might restrict centromeric chromatin, we chose to investigate its role in H3T3P regulation. During mitosis, H3T3P is initially detected chromosome-wide, only to be later targeted to centromeres by the phosphatase activity of the CDCA2-PP1 complex, where CDCA2 (also known as Repo-Man) is essential for targeting of the PP1 phosphatase^30,31^. While the meiotic role of CDCA2 is unknown, it appeared to be highly transcribed at the meiosis-I to spermiogenesis transition^8^. We therefore monitored CDCA2 localization in metaphase-I spermatocytes by IF. Control metaphase-I spermatocytes exhibited both chromatin associated and cytosolic pools of CDCA2 (Fig. 3c, row 1). In the absence of ARID2, chromatin bound CDCA2 was frequently undetectable (51%) (Fig. 3c, row 2). The remainder of the *Arid2^cKO^* spermatocytes exhibited either a reduced (4%) or diffuse (26%) CDCA2 targeting to chromatin (Fig. 3c, row 3,4). Interestingly, these defects in CDCA2 targeting appeared consistent with the changes in H3T3P distribution observed in ARID2 deficient metaphase-I spermatocytes (Fig. 3a, c). Therefore, ARID2 might constrain H3T3P by regulating the chromosomal targeting of CDCA2.

To further investigate this mechanism, we examined BRG1 interacting proteins that were previously identified by immunoprecipitation-mass spectrometry (IP-MS)^5^. While CDCA2 was absent from this list of candidates, we noticed an association between BRG1 and Protein phosphatase 2 regulatory subunit A alpha (PPP2R1A), a scaffolding subunit of PP2A, whose phosphatase activity is essential for the chromosomal targeting of CDCA2^32^ (Extended Fig. 4a; left). ARID2 and BRG1 co-immunoprecipitations (Co-IP) confirmed the association with PPP2R1A (Fig. 3d; lane 1-2, Extended Fig. 4a; right panel, lane 1). Reverse pulldowns further validated the association with BRG1 but failed to reveal an interaction with ARID2 (Fig. 3d; lane 3, Extended Fig. 4a; right, lane 4). This is likely due to the transient nature of the interaction combined with lower expression of *Arid2* relative to *Brg1* (Fig. 1a, Extended Fig. 4a; left). In addition to PPP2R1A, we also detected PP2A, catalytic subunit, PPP2CA^33^ in the ARID2 Co-IP (Extended Fig. 4a; right, lane 1). These associations prompted us to examine PPP2R1A and PPP2CA localization in metaphase-I spermatocytes in response to the loss of ARID2. While PPP2R1A was detected on chromatin and cytosolically in control metaphase-I spermatocytes, it appeared depleted in majority of the mutant metaphase-I spermatocytes (PLK1^-^, 71%) (Fig. 3e). Consistent with their interaction, PPP2CA was seen overlapping ARID2 in control metaphase-I spermatocytes. In contrast, PPP2CA appeared diminished with diffuse localization or was absent in *Arid2^cKO^* metaphase-I spermatocytes (Extended Fig. 4b). Therefore, the destabilization of the PP2A complex probably explains the chromosome-wide spreading of H3T3P in the absence of ARID2. In fact, the majority of *Arid2^cKO^* metaphase-I spermatocytes with reduced PPP2CA levels (83%) displayed H3T3P spreading (Fig. 3f). We conclude that ARID2 impedes H3T3P spreading during metaphase-I by facilitating PP2A mediated targeting of CDCA2 to meiotic bivalents.

Both H3T3P and H2AT120P are required for the recruitment of the chromosome passenger complex (CPC)^4,27,31,34,35^, a process necessary for chromosome segregation. We therefore wanted to determine the impact of displaced H3T3P and H2AT120P on CPC localization in *Arid2^cKO^* metaphase-I spermatocytes. For this purpose we chose CPC factors known to participate in meiosis, such as Inner centromere protein (INCENP), Aurora kinase C (AURKC, meiosis specific) and Aurora kinase B (AURKB)^36,39^. We first monitored phosphorylated INCENP (pINCENP), considered a member of the CPC localization module^40^. Similar to previous studies^39^, pINCENP was detected at centromeres and along the ICA in normal metaphase-I spermatocytes and internal controls from *Arid2^cKO^* testis. In contrast, pINCENP was displaced from ICA and centromeres, adopting a predominantly cytosolic localization pattern in mutant metaphase-I spermatocytes (87%) (Fig. 4a). Similar to pINCENP, its meiotic kinase, AURKC was also depleted from chromatin and enriched cytosolically in mutants (70%) compared to internal controls and normal metaphase-I spermatocytes (Fig. 4b). Since AURKC is known to be partially compensated by AURKB during female meiosis^36^, we were curious to examine AURKB localization in the absence of ARID2. In normal metaphase-I spermatocytes, AURKB was restricted to chromosomes, displaying prominent centromeric enrichment along with diffuse chromatid localization. In contrast, we observed a striking loss of chromosomal AURKB in ARID2 deficient metaphase-I spermatocytes (82%) (Fig. 4c). Therefore, CPC localization is severely impacted in the absence of ARID2. We hypothesize that the mis-regulation of centromeric chromatin contributes to the defect in CPC recruitment.

**Figure 4.**
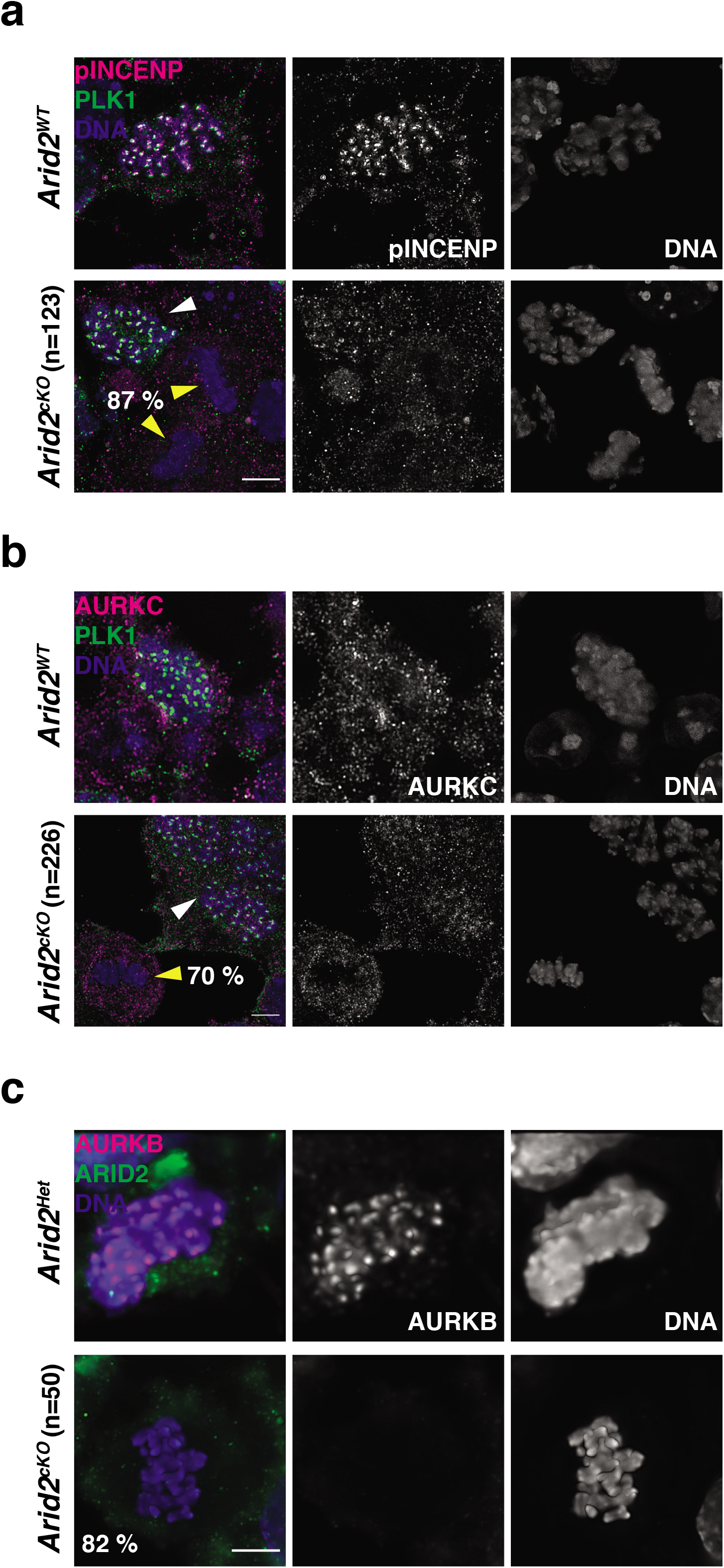
ARID2 influences the localization of the meiotic chromosome passenger complex. (a,b) *Arid2^WT^* and *Arid2^cKO^* metaphase-I spermatocyte squashes immunolabelled for PLK1 (green) and (a) pINCENP (magenta), (b) AURKC (magenta). Internal controls (white arrowhead) and mutants (yellow arrowhead) are indicated. (c) *Arid2^Het^* and *Arid2^cKO^* metaphase-I spermatocyte squashes immunolabelled for AURKB (magenta) and ARID2 (green). DNA stained with DAPI. (a-c) Total number of metaphase-I spermatocytes scored (n) from *Arid2^cKO^* seminiferous tubules and proportion (%) of mutant cells are indicated. Scale bar: 5 μm, magnification: 100.8x.

Our studies highlight a crucial requirement for ARID2 and therefore the PBAF complex in reductional male meiosis. This adds to previously identified roles of SWI/SNF in spermatogonial stem cell maintenance and meiotic prophase-I progression^5^. We show that ARID2 is essential for (i) normal spindle assembly, (ii) PLK1 association with centromeres, (iii) maintaining centromere identity and (iv) the chromosomal targeting of the CPC during metaphase-I, activities that normally promote metaphase exit (Fig. 5). Therefore, it is reasonable to assume that PBAF might function to alleviate the SAC in normal metaphase-I spermatocytes. Interestingly, the loss of ARID2 mimics spindle poisons, that inhibit microtubule synthesis and activate a SAC induced metaphase arrest. In contrast to its meiotic role, the PBAF complex prevents aneuploidy during mitosis^41^, implying a role in SAC activation. These distinct roles might be attributed to differences in mitotic and meiotic PBAF associations. In spermatocytes we demonstrate that ARID2 interacts with PLK1, a kinetochore associated protein, known to regulate anaphase onset during mitosis^42^ and oocyte meiosis^17^. Therefore, ARID2 might mitigate SAC by facilitating PLK1 recruitment to centromeres. Additionally, the resolution of unattached kinetochores is necessary to overcome SAC. Here, the role of ARID2 in spindle assembly is instructive. Interestingly, ARID2 is abundantly detected at spindle poles in metaphase-I spermatocytes (Extended Fig. 2b). These sites usually represent centrosomes, a microtubule organizing center, known to be present in spermatocytes^43^. It is intriguing to speculate whether ARID2 regulates spindle assembly by promoting centrosome activity. Alternatively, known mechanisms of spindle formation involving PLK1 and the CPC^18–20–44^, whose localization are dependent on ARID2 in metaphase-I spermatocytes cannot be ruled out.

**Figure 5.**
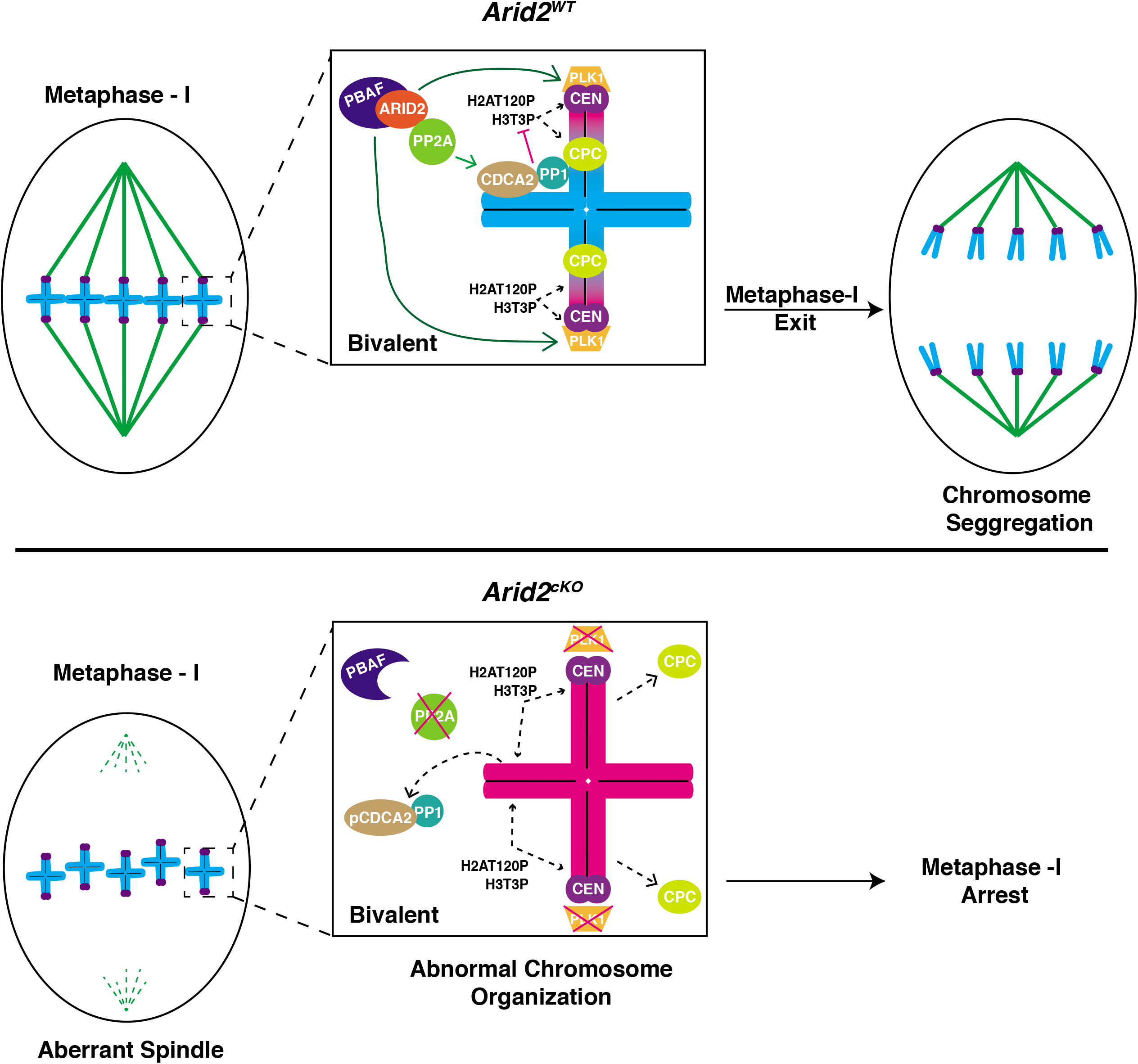
Model describing the role of ARID2 in spermatocyte cell division. ARID2 is required for normal spindle assembly and chromosome segregation in metaphase-I spermatocytes. The association with essential cell division factors such as PLK1 and PP2A implicates PBAF in the regulation of metaphase-I exit. In the absence of ARID2, spermatocytes arrest at metaphase-I, characterized by the lack of spindle, absence of kinetochore associated PLK1, chromosome-wide distribution of H3T3P and H2AT120P and mis-localization of meiotic chromosome passenger complex (CPC). The mis-targeting of H3T3P in the absence of ARID2 occurs due to destabilization of PP2A which results in the reduced chromosomal occupancy of H3T3P phosphatase, CDCA2-PP1. CEN: Centromere

In addition to spindle assembly, PBAF also limits H3T3P and H2AT120P in spermatocytes. In the case of H3T3P, ARID2 appears to employ a conserved mitotic mechanism involving the PP2A regulated chromosomal targeting of CDCA2-PP1, a known H3T3P phosphatase^31,32^. The fact that this pathway is governed by an association between PBAF and PP2A scaffolding subunit, PPP2R1A is novel. Distinct mechanisms might govern the regulation of H2A120P by ARID2, given that H2AT120P is thought to precede H3T3P. In this regard, the mitotic role of Remodeling and spacing factor1 (RSF1), a member of the ISWI family of ATPases, is informative. Like ARID2, RSF1 is required for the centromeric deposition of PLK1 ^45^ and promotes centromeric H2AT120P^46^. The latter activity is achieved by suppressing H2A acetylation.

Given that H3T3P and H2AT120P are known to recruit CPC, the eviction of CPC from chromatin featuring ectopic H3T3P and H2AT120P was interesting. Similar defects in the chromatin association of AURKC in oocytes and TFIID during mitosis in response to an excess of H3T3P have been previously reported ^27,47^. Whether these perturbations result from reduced chromatin accessibility remain unknown. Interestingly, ARID2 prevents the spreading of heterochromatic H3K9me3 in metaphase-I spermatocytes, suggesting a role in promoting chromatin accessibility.

## Acknowledgements

We thank Magnuson lab members for helpful comments on manuscript preparation. We thank Dr. Jesse Raab (University of North Carolina, Chapel Hill) for sharing the *Arid2* floxed mice.

We thank Dr. Michael Lampson (University of Pennsylvania) and Dr. Atilla Töth (Technishe Universität Dresden) for generously providing antibodies. This work was supported by National Institutes of Health grants R01GM101974.

## Author contributions

D.U.M and T.M conceptualized and designed the project. Data curation and validation done by D.U.M. Writing was performed by D.U.M, reviewed and edited by T.M, D.U.M. Project supervision and funding acquisition done by T.M.

## Materials and Methods

### Generation of *Arid2* conditional deletion and genotyping

A knockout first allele of *Arid2* (*Arid2^tm1a(EUCOMM)Wtsi^*) was obtained from EUCOMM^48^ and the mice were generated at the regional mutant mouse resource and research center (MMRRC; Stock number: 036982-UNC; www.mmrrc.org) at the university of North Carolina at Chapel Hill. The lacZ gene trap was flipped out to generate *Arid2 floxed* mice (*Arid2^tm1c(EUCOMM)Wtsi^*) that were maintained on a mixed genetic background. Conditional knockout was achieved using *Stra8-Cre* (activated only in males at P3) ^13^. *Arid2^fl/fl^* females were crossed to *Arid2^fl/+^; Stra8-Cre^Tg/0^* males to obtain *Arid21^flΔ^;Stra8-Cre^Tg/0^* (*Arid2^cKO^*), *Arid21^flΔ^;Stra8-Cre^Tg/0^* (*Arid2^Het^*) and *Arid2^fl+^* (*Arid2^WT^*) males. Genotyping primers used in this study include: *Arid2^fl+^* alleles - (F)- 5’ - CTGCTTAGCCCAAAGGTGTC −3’; (R) 5’ - GACAGTGACTTCAGCTGACC −3’, the excised allele (Δ)- forward primer used above in combination with (R) 5’ -CTGAGCCCAGGTGTTTTTGT-3’ and *Stra8-Cre* - (F) 5’ -GTGCAAGCTGAACAACAGGA-3’, (R) 5’ -AGGGACACAGCATTGGAGTC-3’. All animal work was carried out in accordance with approved IACUC protocols at the University of North Carolina at Chapel Hill.

### Histology

Testes and cauda epididymides from adult *Arid2^WT^* and *Arid2^cKO^* mice were fixed and stained as described previously^49^. Periodic acid-Schiff (PAS), hematoxylin and eosin staining, and sectioning of tissue was performed by the animal histopathology core at the University of North Carolina at Chapel Hill. Seminiferous tubule staging and cell type identification were based on methods outlined previously^50,51^.

### Immunofluorescence staining

Immunofluorescence (IF) was conducted on testes cryosections, squashes and spermatocyte spreads obtained from juvenile (P22 - P27) and adult *Arid2^WT^, Arid2^Het^* and *Arid2^cKO^* mice. IF for metaphase spermatocytes were performed on both testes cryosections and seminiferous tubule squashes. We ensured that IF results were consistent across preparations. Testis cryosections (8-10 μm thick) were prepared and immunostained as previously described^5^. Identical method was used to immunolabel seminiferous tubule squashes, prepared by previously described technique^52^ with one minor modification. To ensure that the squashes adhered to the surface of the glass slide prior to coverslip removal, inverted slides were subjected to downward pressure for 20 minutes by placing above them an eppendorf rack bearing added weight in the form of an aluminum heating block or a filled 500 ml glass bottle. “3D-preserved” prophase-I spermatocyte spreads were prepared and immunolabeled using previously described methods^5,25^. Entire list of primary and secondary antibodies used for IF are provided in supplemental information (Extended table. 1). Images were acquired using Zeiss AxioImager-M2 and Leica Dmi8 fluorescent microscopes. Z-stacks were deconvoluted by Axiovision (AxioImager images) or Huygens (Leica Dmi8 images) software. Image processing and pixel quantification were performed with Fiji^53^.

### Preparation of metaphase spreads

DAPI stained metaphase chromosome spreads were prepared from P19 - P27 and adult *Arid2^WT^, Arid2^Het^* and *Arid2^cKO^* testes, using a previously described technique^54^.

### Preparation of nuclear lysates for immunoprecipitation

Nuclear lysates were obtained from spermatocyte enriched populations isolated from 3-week-old males by methods described previously^55,56^. Protein extracts for immunoprecipitation were isolated from nuclei using a high salt extraction method^5^.

### Co-Immunoprecipitation (Co-IP)

Co-IP were performed exactly as described previously^5^ with minor modifications. These include (i) Incubation of antibodies with nuclear lysates (500 - 600 μg/IP) prior to capture of antibody-antigen complexes with magnetic protein A beads for 3 hours over a rotator at 4 C, (ii) Milder IP wash conditions involving two IP buffer washes, followed by sequential washes once in high salt wash buffer, low salt wash buffer and final wash buffer.

### Preparation of acid extracted histones

Histones were extracted from spermatogenic cells obtained from P23 and P27 *Arid2^WT^, Arid2^Het^* and *Arid2^cKO^* mice using acid extraction protocol described previously ^57^.

### Western blotting

Protein samples were separated by polyacrylamide gel electrophoresis and immunoblotted as described previously^5^. Sample loading was assessed by staining blots with REVERT™ total protein reagents (LI-COR). Blots were scanned on a LI-COR Odyssey imager. All the antibodies used in this study and their corresponding dilutions are listed (Extended table. 1).

### Data sets analyzed

RNA-seq data from purified male germ cells was previously generated (GEO accession: GSE35005)^8^. PPP2R1A association with SWI/SNF was obtained from unpublished data generated using BRG1 IP-MS reported previously^5^.

## Extended Data

### Extended figure legends

**Extended figure 1.**
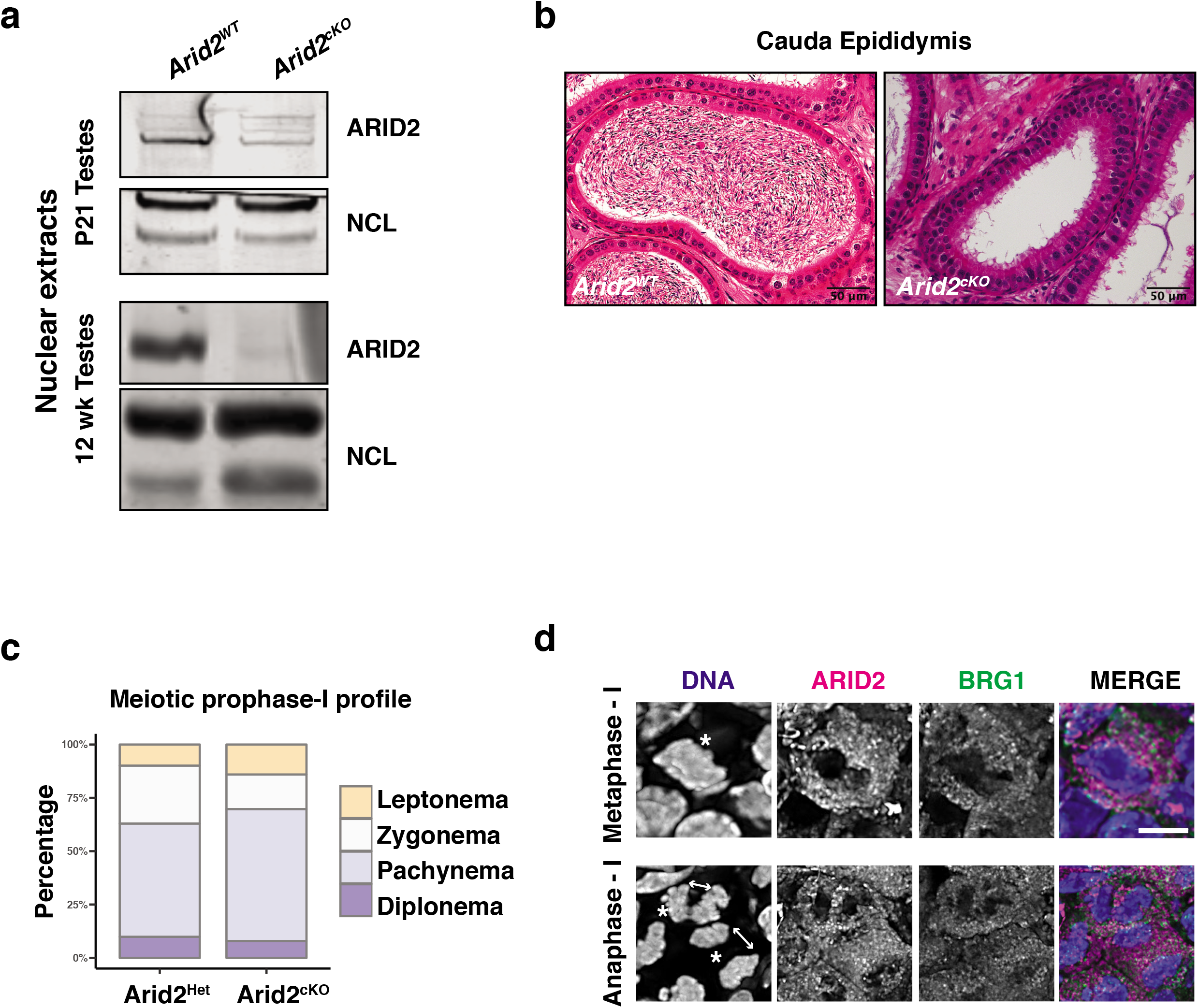
Characterization of ARID2 during spermatogenesis. (a) Western blot displaying ARID2 knockout in spermatogenic cells obtained from juvenile (P21), adult (12-week) *Arid2^WT^* and *Arid2^cKO^* testes. Nucleolin (NCL) was used as nuclear loading control. (b) H&E stained paraffin sections of cauda epididymides obtained from adult *Arid2^WT^ and Arid2^cKO^* mice. Scale bar: 50 μm, magnification: 25x. (c) Meiotic prophase-I profiles obtained from juvenile *Arid2^Het^ and Arid2^cKO^* spermatocyte spreads. Prophase-I substages were identified by SYCP3 and γH2Ax immunostaining. (d) ARID2 (magenta) and BRG1 (green) localization in metaphase-I and anaphase-I spermatocytes. DNA counterstained with DAPI. Scale bar: 5 μm, magnification: 100x.

**Extended figure 2.**
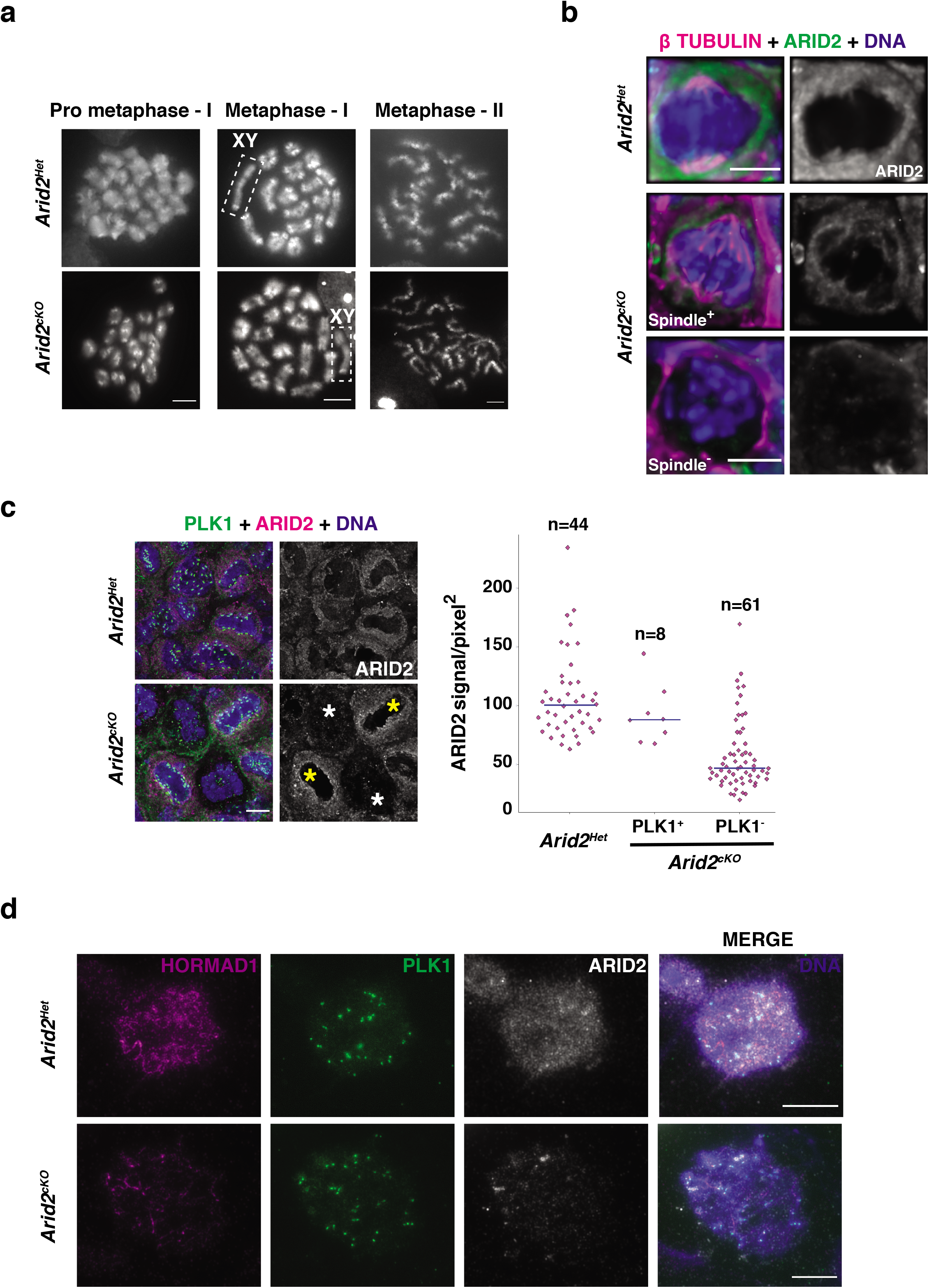
Features of *Arid2^cKO^* spermatocytes. (a) DAPI-stained *Arid2^Het^* and *Arid2^cKO^* metaphase spreads, scale bar: 5 μm, magnification:100x. (b, c) Metaphase-I spermatocytes from *Arid2^Het^* and *Arid2^cKO^* testes cryosections immunolabelled for (a) β-Tubulin (magenta) and ARID2 (green), (c) PLK1 (green) and ARID2 (magenta) with quantification (right) of ARID2 signal (y-axis) from *Arid2^Het^ and Arid2^cKO^* internal control (yellow asterisk, PLK1^+^) and mutant (white asterisk, PLK1) metaphase-I spermatocytes. Number of spermatocytes quantified under each category (x-axis) are indicated (n). (d) *Arid2^Het^ and Arid2^cKO^* diplotene spermatocytes immunolabeled for HORMAD1 (magenta), PLK1 (green) and ARID2 (white). All immunostained preparations were counterstained with DNA. Scale bars: 5 μm, magnification: 100.8x

**Extended figure 3.**
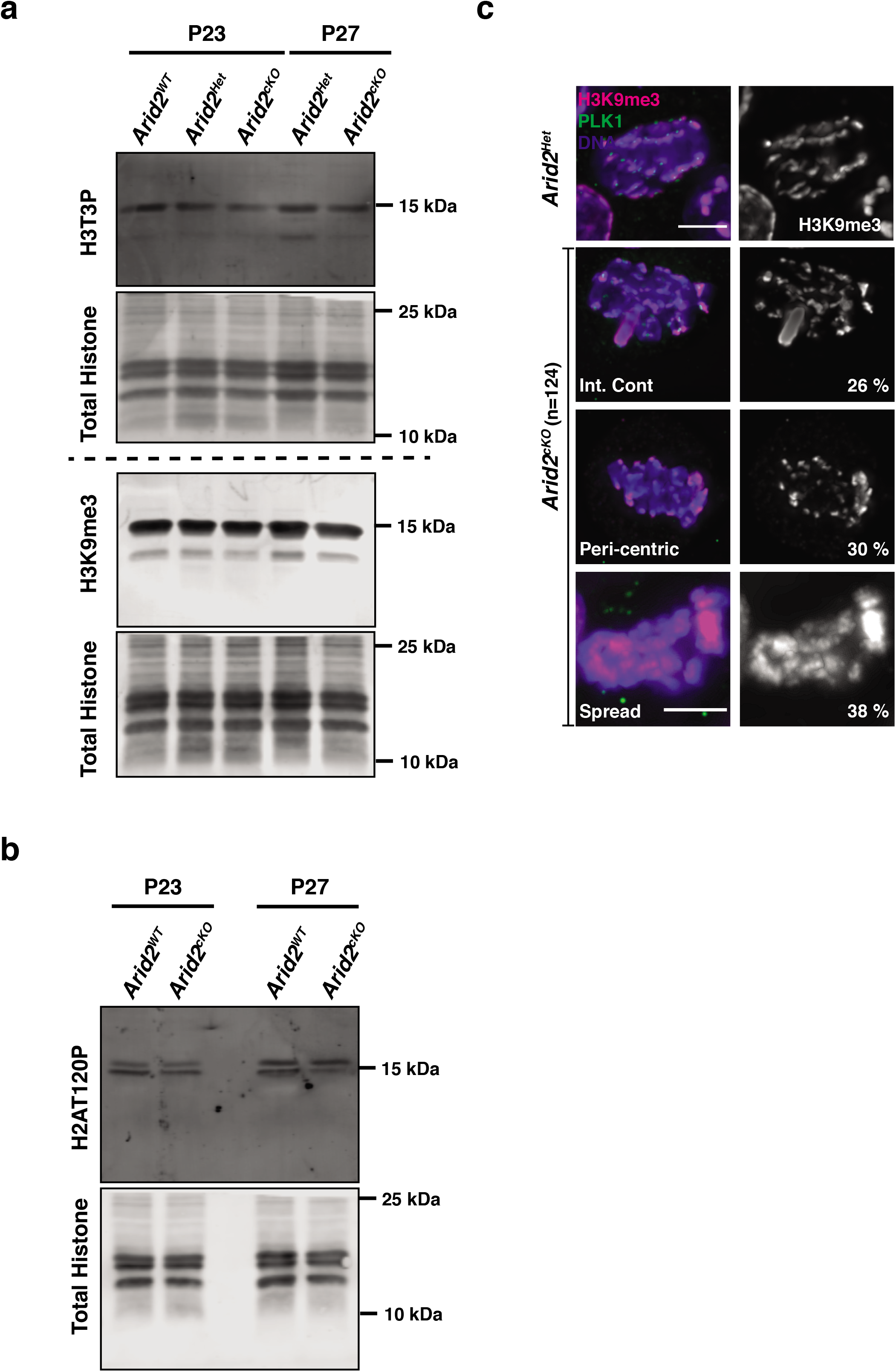
Effect of ARID2 on centromeric and pericentric histone modifications. (a, b) Western blots on acid extracted histones obtained from P23, P27 control and *Arid2^cKO^* spermatogenic cells, depicting abundance of (a) H3T3P (top), H3K9me3 (bottom), and (b) H2AT120P. Total protein from each sample are displayed. (c) Control and *Arid2^cKO^* metaphase-I spermatocytes immunolabelled for H3K9me2 (magenta) and PLK1 (green). DNA stained with DAPI. Total number of metaphase-I spermatocytes scored from *Arid2^cKO^* seminiferous tubules are indicated (n). Scale bar: 5 μm, magnification: 100.8x.

**Extended figure 4.**
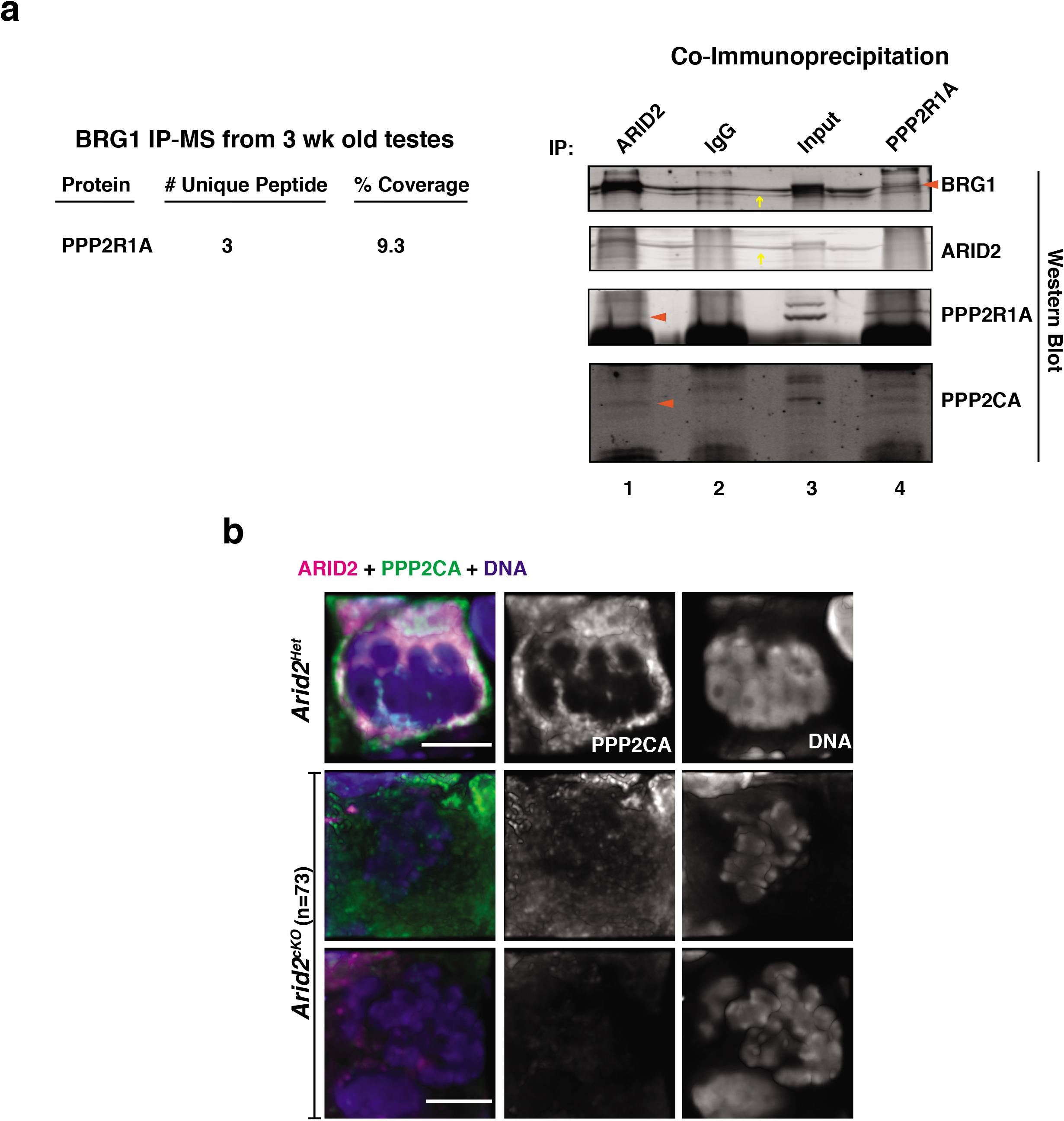
ARID2 associates with PP2A. (a) Table (left) summarizing PPP2R1A interaction with BRG1 previously identified by IP-MS and ARID2, PPP2R1A Co-IP’s (right). Red arrowheads indicate interacting proteins, yellow arrow denotes non-specific streak. Numbers at bottom denote sample lanes. (b) Control and *Arid2^cKO^* metaphase-I spermatocytes immunolabelled for ARID2 (magenta) and PPP2CA (green). Total number of metaphase-I spermatocytes scored from *Arid2^cKO^* seminiferous tubules are indicated (n). Scale bar: 5 μm, magnification: 100.8x

**Extended table 1.** List of antibodies

| Antibody | Vendor | Application |
| --- | --- | --- |
| Rabbit anti-ARID2 | Thermofisher (PA5-35857) | IF (1:100)<br>IP (10 µg)<br>WB (1:1000) |
| Mouse anti- BRG1 (G-7) | SantaCruz Biotech (sc- 17796) | IF (1:100)<br>WB (1:1000) |
| Rabbit anti- BRG1 | Abcam (ab110641) | IP (7 µg)<br>IF (1:500) |
| Mouse anti- γH2Ax | Millipore (05-636) | IF (1:1000) |
| Rabbit anti NCL (Nucleolin) | Bethyl (A300-711A) | WB (1:5000) |
| Mouse anti- SYCP3 | Abcam (ab97672) | IF (1:500) |
| Rabbit anti- γH2Ax | Cell signaling (9718) | IF (1:1000) |
| Rabbit anti- H3S10P | Millipore (06-570) | IF (1:500) |
| Mouse anti β-TUBULIN | DHSB (E7) | IF (1:25) |
| Rabbit anti- CENPA | Abcam (ab33565) | IF (1:25) |
| Mouse anti- PLK1[35-206] | Abcam (ab17056) | IF (1:200)<br>WB (1:1000) |
| Guinea pig anti- HORMAD1 | Gift from Dr. Atilla Töth, TU Dresden | IF (1:500) |
| Rabbit anti- H3T3P | Active Motif (39153) | IF (1:100)<br>WB (1:1000) |
| Rabbit anti- H2AT120P | Active Motif (39392) | IF (1:100)<br>WB (1:1000) |
| Rabbit anti- H3K9me3 | Abcam (ab8898) | IF (1:500)<br>WB (1:1000) |
| Rabbit anti- CDCA2 | Sigma (HPA030049) | IF (1:100) |
| Rabbit anti- PPP2R1A | Proteintech (15882-1-AP) | IF (1:100)<br>IP (1.6 µg – 8 µg)<br>WB (1:800) |
| Mouse anti- PPP2CA | DHSB (CPTC-PP2A-4) | IF (1:10)<br>WB (1:20) |
| Rabbit anti- phosphoINCENP | Gift from Dr. Michael Lampson, University of Pennsylvania | IF (1:500) |
| Rabbit anti – AURKC | Bethyl (A300-BL1217) | IF (1:30) |
| Mouse anti- AURKB | BD biosciences (611082) | IF (1:30) |
| **Secondary antibodies** |  |  |
| Goat anti-mouse, Alexa fluor 488 | ThermoFisher (A-11029) | IF (1:500) |
| Goat anti-mouse, Alexa fluor 568 | ThermoFisher (A-11031) | IF (1:500) |
| Goat anti-rabbit, Alexa fluor 568 | ThermoFisher (A-11036) | IF (1:500) |
| Goat anti- rabbit, Alexa fluor 488 | ThermoFisher (A-11008) | IF (1:500) |
| Goat anti- guinea pig, Alexa fluor 568 | ThermoFisher (A-11075) | IF (1:500) |
| Goat anti- rabbit, Alexa fluor 647 | ThermoFisher (A-21245) | IF (1:500) |
| IRDye 680LT Goat anti-rabbit | LI-COR (926-68021) | WB (1:20,000) |
| IRDye 680LT Goat anti-mouse | LI-COR (925-68020) | WB (1:20,000) |
| IRDye 800CW Goat anti-mouse | LI-COR (926-32210) | WB (1:10,000) |

